# COMPUTATIONAL IDENTIFICATION OF ANTIBODY-BINDING EPITOPES FROM MIMOTOPE DATASETS

**DOI:** 10.1101/2023.12.17.572067

**Authors:** Rang Li, Sabrina Wilderotter, Madison Stoddard, Debra Van Egeren, Arijit Chakravarty, Diane Joseph-McCarthy

## Abstract

A fundamental challenge in computational vaccinology is that most B-cell epitopes are conformational and therefore hard to predict from sequence alone. Another significant challenge is that a great deal of the amino acid sequence of a viral surface protein might not in fact be antigenic. Thus, identifying the regions of a protein that are most promising for vaccine design based on the degree of surface exposure may not lead to a clinically relevant immune response. Linear peptides selected by phage display experiments that have high affinity to the monoclonal antibody of interest (“mimotopes”) usually have similar physicochemical properties to the antigen epitope corresponding to that antibody. The sequences of these linear peptides can be used to find possible epitopes on the surface of the antigen structure or a homology model of the antigen in the absence of an antigen-antibody complex structure. Herein we describe two novel methods for mapping mimotopes to epitopes. The first is an ensemble approach, which combines the prediction results from two existing methods. The second is a novel algorithm named MimoTree, that allows for gaps in the mimotopes and epitopes on the antigen. More specifically, a mimotope may have a gap that does not match to the epitope to allow it to adopt a conformation relevant for binding to an antibody and residues may similarly be discontinuous in conformational epitopes. MimoTree is a fully automated epitope detection algorithm suitable for the identification of conformational as well as linear epitopes.

## 1 Introduction

Antibodies are produced by B cells during the immune response to the presence of foreign matter within the body (“antigens”). Antibodies recognize and bind to specific regions of target antigens to perform their functions, and the regions of antigen molecules to which antibodies attach are defined as epitopes (Sela-Culang et al., 2013). B-cell epitopes can be classified into linear and conformational epitopes; in the latter case, epitopes consist of patches of residues that lie close to each other in three-dimensional space but are separated in amino acid sequence (Potocnakova et al., 2016, Sanchez-Trincado et al., 2017). Conformational epitopes generally account for about 90% of overall antibody binding to an antigen (Najar et al., 2017). In addition, conformational epitopes may offer functional advantages over linear epitopes that confer enhanced viral neutralization (Vinion-Dubiel et al., 2001) and longer lasting immunity (Steimer et al., 1991). For example, conformational epitope-targeting antibodies to the HIVgp120 glycoprotein are more effective at neutralizing viral isolates than linear epitope-targeting antibodies to the same protein, and they are also responsible for the majority of gp120-specific CD4-blocking activity in HIV-1-infected human sera.

A deeper knowledge of the structure and immunogenic properties of conformational epitopes may be beneficial for designing interventions for many therapeutic areas, including viral infections and cancer (Saxena et al., 2006, Tegoni et al., 1999, Li et al., 2005, Nybakken et al., 2005, Adams and Weiner, 2005, Cho et al., 2003, Riemer et al., 2005). In particular, this knowledge is particularly relevant to vaccine design, since it is generally safer yet comparably effective to vaccinate with the epitope only compared to the inactivated antigen (De Groot, 2004). Immune responses elicited by partial antigens may be sufficient to provide competent protection since most of the surface structure of the antigen molecules are not antigenic (Sanchez-Trincado et al., 2017). Therefore, identifying immunogenic conformational epitopes and developing strategies to generate an effective immune response against the identified epitopes are important problems in computational vaccinology.

Mimotopes represent a solution to the challenges above. A mimotope is a peptide that mimics the structure of an epitope, causing an antibody response that is similar to the one elicited by the epitope (Sidhu et al., 2003). Thus, an antibody for a given antigen will recognize the mimotopes which mimic the corresponding epitope since antibodies usually do not distinguish between epitopes and epitope-based proteins (Matsubara, 2022). Mimotopes are commonly obtained from phage display libraries through bio-panning experiments (Moreau et al., 2006, Bublil et al., 2007). Bacteriophages displaying potential mimotope peptides are incubated with the target antibody, which is immobilized on a solid support. As such, specific phages in the library bind to the target antibody. Unbound phages are washed out, while bound phages are selected, eluted, and amplified (Bazan et al., 2012). Multiple rounds of evolution may be performed. This method is intended to select peptides with high binding affinity to the chosen antibodies. A portion of the selected mimotopes will have some homology to antigen epitopes and may trigger the anticipated immune response. As a result, mimotopes can be employed in and potentially used as a starting point for vaccine design.

Mimotopes are especially useful in vaccine design when the epitope is known, as this supports a rational approach. For example, if the epitopes corresponding to the mimotope are known, this allows selection of mimotopes with the best reactivity across antigenic variation, which is important for designing vaccines against rapidly evolving pathogens. The algorithmic task of the mimotope-to-epitope mapping is to map the mimotope to the part of the antigen surface that binds the antibody used to create the mimotope. This epitope should have similar physicochemical properties to the mimotope, facilitating antibody binding to both. Since the mimotope is usually not identical but similar to its corresponding epitope at the sequence and structural level, the algorithm considers residues with a physicochemical property distance within a certain range as identical, which is a major challenge in the mapping process. The regions of the antigen surface that are aligned with the mimotopes have a higher probability of being antigenic. Similar to phage-display experiments, molecular docking has been used to enrich potential target regions of the antigen. Protein-protein docking methods have been used to identify the antibody binding site (the epitope) on an antigen protein (Desta et al., 2022); this approach requires a structure or model of both the antigen and antibody structure. Here we are mapping the mimotope to the surface of the antigen to identify the epitopes that the antibody is expected to be capable of binding. As such, only the structure or model of the antigen protein is required. That is, the process of mapping mimotopes (peptides with high binding affinity to the antibody) to the surface of the macromolecule antigen identifies antibody binding sites on the antigen.

Several computational methods for mimotope mapping based on the antigen structure alone exist (Bublil et al., 2007, Mayrose et al., 2007, Negi and Braun, 2009), all of which have been validated on a similar set of mimotopes and corresponding protein epitopes based on the existence of the corresponding antigen-antibody complex structures. While the overall precision of the predictions is reasonable, sensitivity is low, meaning that residues that lie within the true epitopes are left out of the epitopes predicted by these algorithms. Thus, the existing methods are useful but imperfect.

In addition to broadly low sensitivity, specific mimotope-to-epitope mapping algorithms display unique weaknesses. For example, Mapitope (Bublil et al., 2007) is an algorithm that breaks each mimotope in the dataset into overlapping residue pairs (each residue pair contains two consecutive amino acids in the mimotope sequence), and then computationally pools these residue pairs and ranks the occurrence of each type. Next, it uses the high-frequency occurrences to search on the antigen surface to find the antigen surface residues mapped by these pairs. However, this algorithm requires many statistically relevant parameters customized by the user, and the length of the residue pairs it considers is so short that different parameter settings can have a very large impact on the prediction.

Pupko and co-workers also developed PepSurf (Mayrose et al., 2007), which uses a color-coding algorithm to find all possible simple paths on the antigen surface, matches these paths with mimotopes based on amino acid similarity, and finally clusters the paths with high similarities to get the final prediction. However, the run time of PepSurf depends linearly on the length of the mimotopes because the step of searching for every possible simple path on the antigen surface limits the rate to a great extent. Therefore, PepSurf is only able to process mimotopes with up to 15 amino acids. Subsequently, EpiSearch (Negi and Braun, 2009) solved the run time issue, being able to complete all calculations in less than a minute. However, EpiSearch’s method of dividing regions on the antigen surface is fast but imprecise, and it is also unable to process more than 30 mimotopes at a time.

The overall goal of this study is to develop mimotope-to-epitope mapping prediction methods that are more robust in terms of providing predictions with improved sensitivity. Firstly, we present an ensemble method that combines two existing methods to provide more sensitive predictions. While the mimotope is similar to but usually not identical to the true epitope, most mimotopes produce some gaps (in mimotope and/or epitope sequence) when they are aligned to the antibody surface. In particular, most contiguous mimotope residues correspond to discontinuous binding sites on the antigen surface that bind to the antibody (Cho et al., 2003). To address this feature, we developed a prediction algorithm that takes into account these sequence gaps while also referring back to the three-dimensional (3D) structure of the antigen in the final mimotope mapping step to achieve more sensitive and accurate epitope prediction.

## 2 Methods

### 2.1 Test Set

To assess the performance of the mimotope-to-epitope mapping methods, we collated ten test cases from similar publications (Huang et al., 2008a). The criteria for selecting test cases are i) the 3D crystallography structure of the antigen-antibody is available; ii) the complex contains only the antigen and antibody; iii) a set of mimotopes was derived from bio-panning experiments with the given antibody. The test set is presented in Table 1. Two of the test cases, 1AVZ and 1HX1, are for protein-protein interactions where one protein is considered as the “antigen” here since phage-display libraries of peptides were screened for binding to the other protein considered as the “antibody” in the pair for our purposes. For example, for 1AVZ, a peptide library was screening using phage display to identify peptides that bind to the Fyn SH3 domain (the “antibody”) to block its interactions with negative regulatory factor (Nef) (Rickles et al., 1994, Greenway et al., 1996). The Nef dimer, the biologically relevant form of Nef (Arold et al., 1997, Wu et al., 2018), was taken as the “antigen”.

**Table 1.**
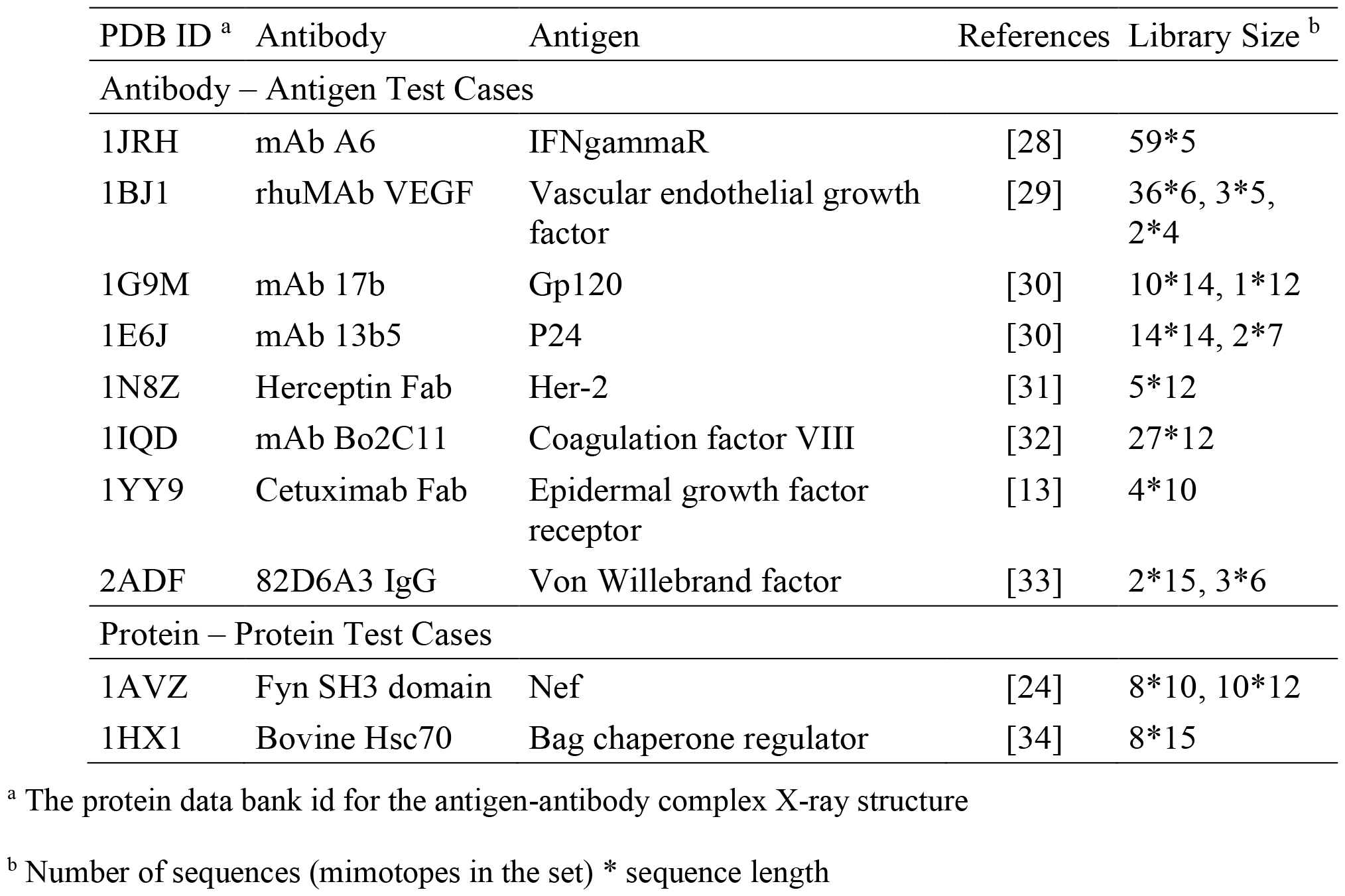
Mimotope-to-Epitope Test Set.

Since mimotope-to-epitope methods are designed to predict the epitope on an antigen surface, as a more stringent test, we also compiled a set of unbound antigen structures when available that correspond to the test set antigen-antibody structures. This is a more rigorous test since the antigen may undergo conformational changes upon binding to the antibody. The antigen-only test set is given in Table 2. For two of the test cases in Table 1, 1N8Z and 1JRH, the corresponding unbound antigen structure does not exist.

**Table 2.**
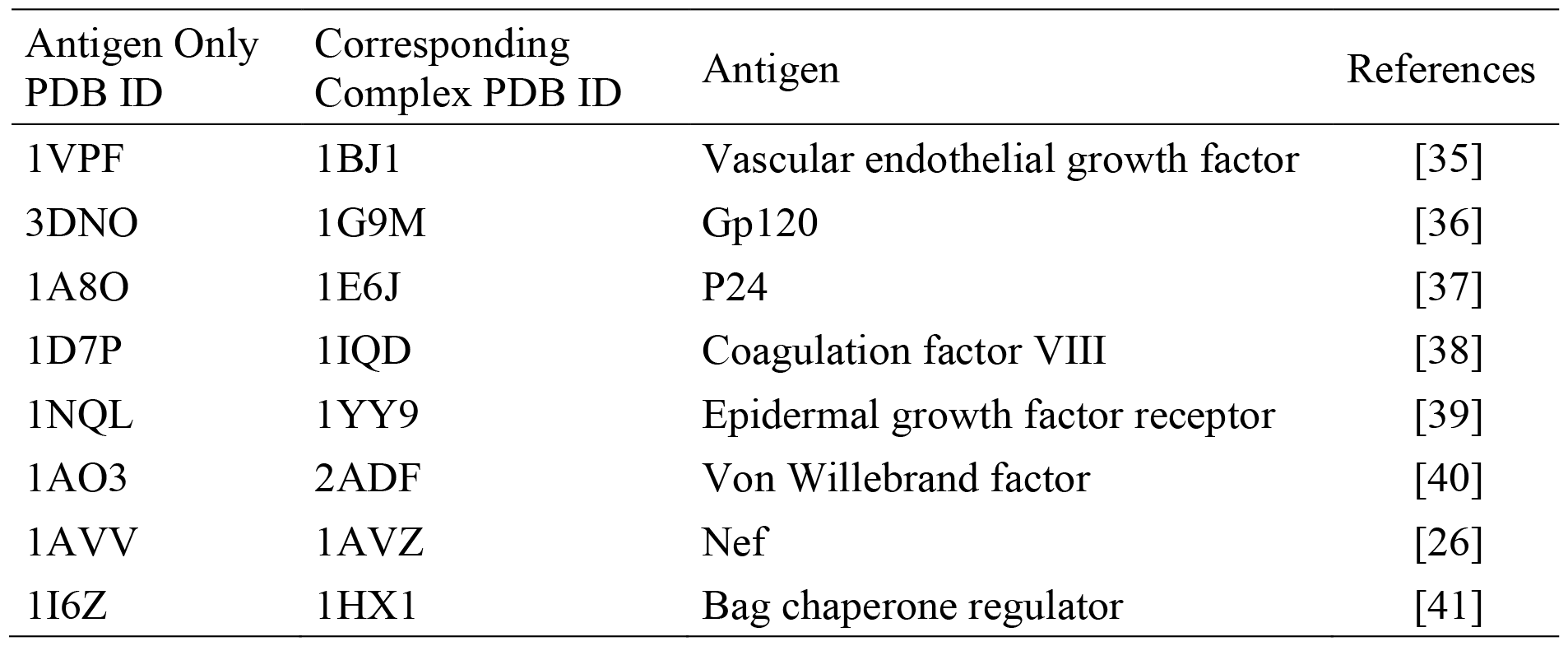
Antigen-Only Structures Corresponding to the Test Set.

| Antigen Only PDB ID | Corresponding Complex PDB ID | Antigen | References |
| --- | --- | --- | --- |
| 1VPF | 1BJ1 | Vascular endothelial growth factor | [35] |
| 3DNO | 1G9M | Gp120 | [36] |
| 1A8O | 1E6J | P24 | [37] |
| 1D7P | 1IQD | Coagulation factor VIII | [38] |
| 1NQL | 1YY9 | Epidermal growth factor receptor | [39] |
| 1AO3 | 2ADF | Von Willebrand factor | [40] |
| 1AVV | 1AVZ | Nef | [26] |
| 1I6Z | 1HX1 | Bag chaperone regulator | [41] |

### 2.2 Epitope Definition

Chimera (Pettersen et al., 2004) was used to define the true epitope for each antigen in the test set. An antigen residue is considered to be part of the epitope if the difference between its solvent-accessible surface area (SASA) in the antigen structure (taken from the complex) and in the antigen-antibody complex is greater than 10 Å^2^ (Hu and Yan, 2009).

### 2.3 Amino Acid Similarity

Since the amino acids in a mimotope are not typically identical to those in the true epitope, the ability of a mimotope to bind with high affinity to a target antibody and possibly achieve the same function as the true epitope is likely due to the fact that at least a portion of the amino acids in the mimotope share similarity rather than identity with the epitope residues. Similarities between amino acids determine whether they can substitute for each other in a sequence while maintaining similar peptide/protein properties and function. Several methods exist for assessing amino acid similarity, based on the physicochemical properties of the individual amino acids, the role they play in the protein structure, or more subtle contributions (Stephenson and Freeland, 2013). We used quantitative descriptors of the properties of amino acids published by Braun and coworkers when performing mimotope to antigen surface mapping (Venkatarajan and Braun, 2001). The calculated similarity is as follows:

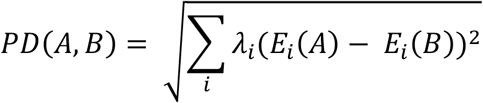

where PD (A, B) represents the property distance between amino acid A and amino acid B, and λ_*i*_ are the eigenvalues corresponding to the eigenvectors E_i_ (i = 1-5) representing the physicochemical properties of each amino acid. The lower the PD between two amino acid types, the more similar they are. According to Braun and co-workers, the finest grouping of amino acids based on the physicochemical properties is defined by a distance cutoff of 9.5 (Venkatarajan and Braun, 2001). In practice, we chose a distance cutoff of 8 based on the run time of the algorithm and accuracy of the output, which means that two amino acids with a PD of 8 or less are considered identical.

### 2.4 Selection of Surface Residues

In order to perform mimotope mapping only on the antigen surface, we defined antigen surface residues as those having a SASA greater than 5% of its maximal SASA (Miller et al., 1987). The SASA value per residue in the antigen structure was calculated using Chimera with a default probe radius of 1.4 Å2 (Pettersen et al., 2004). The maximal SASA of each standard amino acid was defined as the total SASA of the residue in an extended Gly-X-Gly peptide where X represents the residue of interest (Miller et al., 1987).

### 2.5 “The Ensemble Method”

In this study, two existing approaches that have complementary features in their searching algorithms (described below) have been identified, investigated, and combined. The two selected methods are EpiSearch and PepSurf. The final ensemble method takes the union of the top-ranked solutions from EpiSearch (http://curie.utmb.edu/episearch.html) and PepSurf (http://pepitope.tau.ac.il/index.html) calculated using default parameter settings on their respective servers (Negi and Braun, 2009, Mayrose et al., 2007).

#### 2.5.1 Algorithm Description of EpiSearch

EpiSearch first generates surface patches centered on each antigen surface residue using a preset radius value. These overlapping patches can cover the entire antigen surface. The type and number of amino acids contained in each surface patch and in each mimotope sequence are stored in separate matrices. Then, EpiSearch calculates the number of residues in each surface patch that are “identical” to any amino acid in the mimotope, where a pair of residues with a property distance (as defined above) less than or equal to a preset value is defined as a pair of identical residues. For each individual mimotope, EpiSearch scores all surface patches based on the degree of matching of each patch with the mimotope. The highest scoring patch is selected as the predicted patch corresponding to the query mimotope. After obtaining the predicted patches for all the input mimotopes in the set, the individual residues in the predicted patches that match to the corresponding mimotope are predicted to be part of the conformational epitope of the antigen (Negi and Braun, 2009).

#### 2.5.2 Algorithm Description of PepSurf

PepSurf first unfolds the antigen surface onto a surface map containing all surface residues, where a pair of neighboring residues in the map is defined as any two residues within a distance of 4 Å. For each individual mimotope, PepSurf looks for all possible simple paths on the surface map that are the same residue length as the given mimotope. With its color-coding technique, PepSurf is able to find every linear path on the surface map without duplication. Afterward, PepSurf calculates the weight of each path by comparing its similarity to the query mimotope based on a modified BLOSUM62 matrix (Henikoff and Henikoff, 1992, Mayrose et al., 2007). The modified substitution score representing the similarity between amino acids i and j, h (i, j), is calculated as follows:

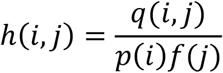

where q(i, j) is the observed probability of occurrence for each i, j pair in the original BLOSUM62 matrix; p(i) and f(j) are the probabilities of occurrence for i and j in the phage display library and in the original BLOSUM62 matrix, respectively (Mayrose et al., 2007). For each mimotope, the surface path with the highest weight is selected. Finally, PepSurf clusters all the selected surface paths and scores them with respect to their similarity to their corresponding mimotopes to obtain the final prediction results for the mimotope set (Mayrose et al., 2007).

#### 2.5.3 Creating an Ensemble Prediction

To integrate EpiSearch and PepSurf, we tested both union and intersection methods for “The Ensemble Method”. First, EpiSearch and PepSurf are used to make predictions for a particular mimotope set-antigen pair independently using default parameters. For EpiSearch, the default patch size which represents the radius value of the surface patches (12 Å), the default PD cutoff (10), and the default accuracy cutoff which indicates the maximum number of mismatches allowed in each surface patch (3) were used. For PepSurf, the default substitution matrix (BLOSUM62) and the default gap penalty (−0.5) were used. The union method is defined as taking the union of the residues in the top-scoring predicted epitope sequences from EpiSearch and PepSurf. The intersection method takes the intersection of the top-scored solutions from EpiSearch and PepSurf. We chose the union method after an initial evaluation the performance of these two methods, which showed that the union significantly improved the sensitivity of the existing methods while intersection decreased performance.

### 2.6 MimoTree

Mimotopes may only partially mimic the structure and function of the true epitopes that they mimic; most mimotopes are similar but not identical to the true epitope both at the level of sequence and structure. Moreover, during the process of the mimotopes binding to the antibody, it is likely that some residues of the mimotopes will not be bound to the antibody residues, but rather will simply exist in the mimotope sequence to allow the mimotope to adopt a conformation relevant for binding to the antibody. As a result, there may be gaps in the mapping of the mimotope sequence to the antigen surface. To address this issue, some existing methods use a gap penalty parameter added to the similarity score of the mimotope and the antigen surface area or to the weighted score of the predicted path, such that the weight of a mimotope that forms gaps is reduced. Introducing a gap penalty, however, reduces the probability that a mimotope with a gap will be highly scored even if it does match a true epitope (Mayrose et al., 2007, Negi and Braun, 2009).

In fact, mimotopes that form gaps during binding should be scored equally well when mapped onto an epitope region. A unique feature of our algorithm is that it allows for these gaps and does not penalize mappings with such gaps. These gaps in the mimotope sequence binding to the antibody are expected to translate into gaps in the mapping of the mimotope onto the antigen surface. Another unique feature of MimoTree is that the 3D structure of the antigen is considered when determining if a gap in the putative epitope mapping on the antigen surface is allowed. That is, if the distance on the antigen surface is within the linear length possible for the number of residues of the mimotope gap, then the mimotope can be mapped at the corresponding position on the surface antigen.

#### 2.6.1 Algorithm Flow

MimoTree is a Depth First Search (DFS)-based algorithm. The inputs to MimoTree are a protein data bank (PDB) file of the structure of the antigen of interest and a mimotope set. MimoTree initially creates a surface map of the input antigen structure. For each individual mimotope in the dataset, MimoTree performs a DFS tree-based search of the mimotope sequence to identify surface segments or seeds on the antigen surface that match to the mimotope. MimoTree then connects seeds on the antigen surface that are within a physically reasonable distance based on the 3D structure of the antigen as described above. The predicted epitope for an individual mimotope is a concatenation of seeds (a seed connection) matched to the antigen surface, where the top three longest seed connections are output as the prediction. The final output of the algorithm is the union of the predictions corresponding to all mimotopes in the mimotope set. A detailed description of the method is outlined in Fig.1.

#### 2.6.2 Creation of a Surface Map

For the antigen, MimoTree creates a dictionary where each surface residue is a key, and a list containing the adjacent surface residues of each key is represented as the corresponding value. The maximum distance at which two residues are treated as adjacent to each other is 4 Å (Madabushi et al., 2002).

#### 2.6.3 Tree-Based Searching to Identify Seeds

For each mimotope in the set, starting from the first amino acid, MimoTree identifies all of the residues on the protein surface that match. If the property distance between two amino acids is within the pre-defined range (described above), the amino acids are considered as matching in this algorithm. For each surface residue that matches the first residue in the mimotope, MimoTree checks the surface map to see if any adjacent residues match the second residue in the mimotope. If so, it continues searching for the next residue in the mimotope sequence. Otherwise, it terminates that search and starts searching in a different direction from that residue. If that fails, the algorithm starts searching from the next antigen surface residue that matches the first mimotope residue. After finding all possible mimotope matching segments on the antigen surface starting from the first residue of the query mimotope, MimoTree starts a new round of surface searching from the second residue of the mimotope. It continues and completes the surface searching starting from each residue in the mimotope. Finally, the output is all the seeds (matching segments) on the antigen surface that match any part of the query mimotope sequence.

This tree-based searching is based on the DFS algorithm. It starts with a matching surface residue, searches from one direction to the end of that path, and then returns to the previous level to start the search from another direction until the end. Each surface search starting from a residue stops only after all possible paths have been searched, ensuring a comprehensive surface search.

#### 2.6.4 Seed Connections

After finding all of the seeds (the residues on the antigen surface that partially match the mimotope), MimoTree tries to join the seeds based on the sequence order of the mimotopes to form “seed connections.” The algorithm connects seeds if the mimotope gap in an extended linear conformation is sufficiently long extend from the one seed to the other seed. That is, the length of the gap in the mimotope sequence, assuming an extended linear conformation, is compared to the distance in the 3D antigen structure from the last residue of the first seed to the first residue of the next seed. If the maximum length of the mimotope gap is greater than or equal to the distance on the 3D antigen structure between the ends of seeds that are being connected, then this mimotope can be mapped at that position (to that epitope) on the antigen surface. In the hypothetical example of a 7-residue mimotope, where residues 1-3 and 6-7 map to the epitope (Fig. 1B and C), the mimotope gap of residues 4-5 is allowed if the length of residues 4-5 plus one residue on either side in an extended linear conformation is greater than or equal to the distance between the residues in the 3D antigen structure that map to mimotope residues 3 and 6. More specifically, the algorithm considers the N-N distance between two adjacent residues in the mimotope as 3.5 Å (Bhagavan and Ha, 2015), so in the example above the maximum distance would be 10.5 Å (3 * 3.5 Å/residue). The N-N distance between the corresponding residues on the surface of the antigen (those that match to mimotope residues 3 and 6) is then calculated from the coordinates of the residues in the input PDB file and must be less than the linear mimotope gap length (10.5 Å in the example) for a connection to be made.

**Figure 1.**
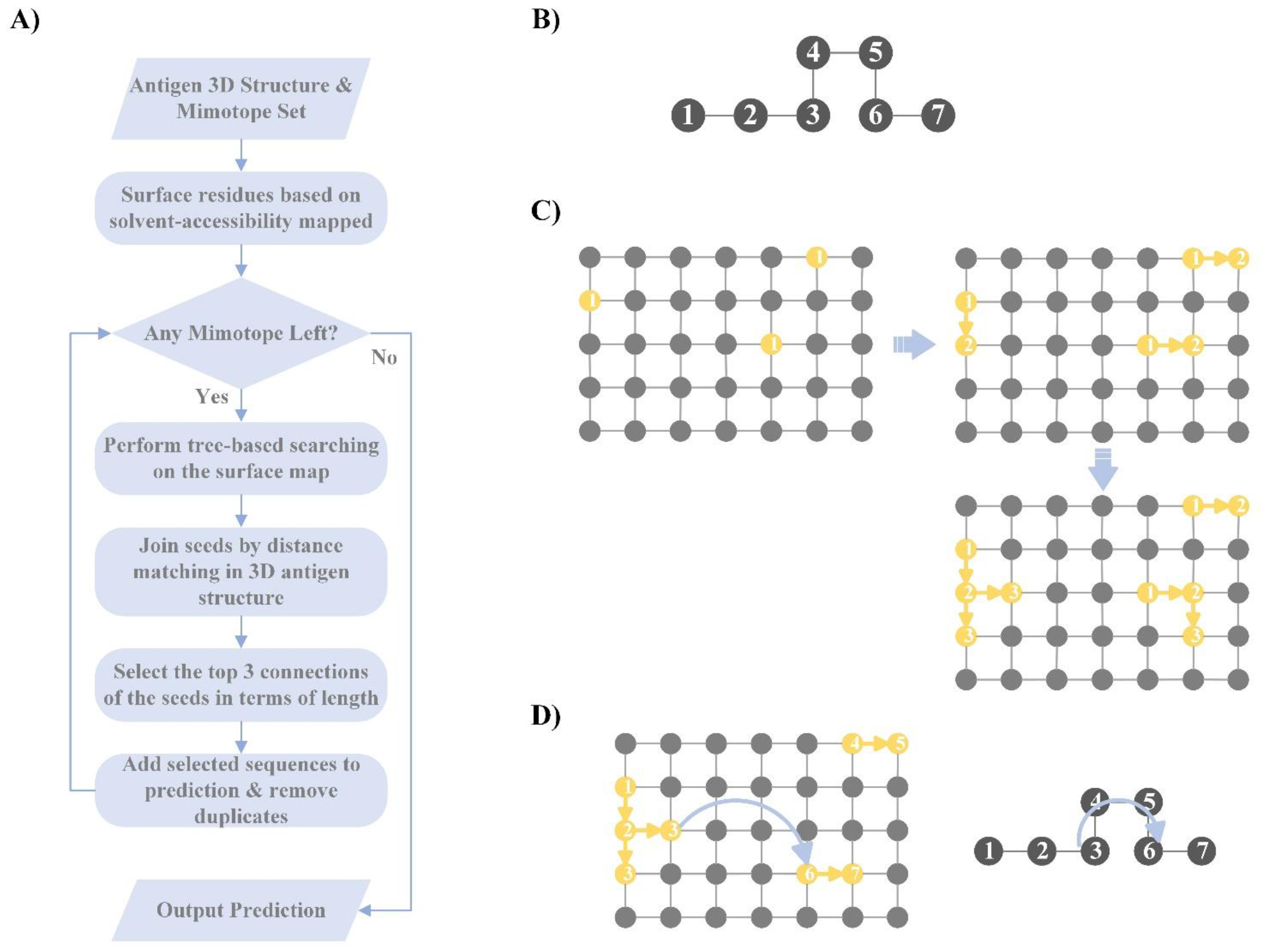
Overview of MimoTree algorithm. A flowchart of MimoTree algorithm is shown in A) and an example of the tree-based search starting from residue 1 of the mimotope sequence given in B) is shown in C).

#### 2.6.5 Selection of the Seed Connections

Since MimoTree searches for possible antigen surface seed connections on the antigen surface based on the sequence of mimotope, the longer the seed connection is, the greater the portion of the mimotope that is matched and the more likely it is that a true epitope has been identified. The length of a seed connection excludes gaps (so is length 7 for the example above) and can extend to the limit of the length of the mimotope (10 for the example above). For each individual mimotope, MimoTree takes the union of the top three longest connections of the seeds to be the predicted epitope of the query mimotope. The final prediction of MimoTree is the union of the predictions corresponding to all the mimotopes in the mimotope set.

#### 2.6.6 Parameter Tuning for MimoTree

MimoTree contains a tunable parameter, the cutoff value for the degree of amino acid similarity. As mentioned previously, we calculate the property distance between any two amino acids using the values of the physicochemical descriptors of each standard amino acid and the eigenvalues indicating the weights of different descriptors. When the property distance between two amino acids is less than a preset cutoff value, MimoTree considers the two amino acids to be identical. Thus, in the process of mapping mimotopes to the antigen surface, if the property distance between a residue in the mimotope and a residue on the antigen surface is less than the cutoff value, the algorithm aligns the mimotope residue to that position on the antigen surface.

If the cutoff value is decreased, the algorithm will match fewer but more similar mimotope residues to antigen residues. As a result, the output will be possible epitopes that are more similar in property space to the aligned mimotopes. Thus, if the input mimotopes are very similar to the true epitope, reducing the cutoff will improve both sensitivity and precision of the algorithm, but if the input mimotopes are not similar enough to the true epitope, lowering the cutoff will reduce both sensitivity and precision. In other words, lowering the cutoff makes the algorithm more dependent on the similarity or identity between the mimotope and the true epitope, which reduces the robustness of the algorithm. Since lowering the threshold would consider fewer residues, it would also shorten the run time of the program.

If the cutoff value is increased, the algorithm will match mimotope residues to some antigen surface residues that are not as similar; this will increase sensitivity because the algorithm will analyze more surface residues, but it will likely decrease precision. However, since residues that are more dissimilar to those in the mimotopes will be included, this move could improve the robustness of the algorithm since some matches will be made even if the input mimotopes are not identical to the true epitope. However, the running time of the program will be increased due to the tolerance of the reduced similarity.

In practice, we found that a cutoff value of 8 yielded the best results across the test set based on the accuracy and run time. All results obtained by MimoTree herein utilize a cutoff of 8.

### 2.7 Incorporation of MimoTree into “The Ensemble Method”

We next sought to add MimoTree into “The Ensemble Method” to evaluate if the performance would be improved relative to MimoTree alone or the ensemble of EpiSearch and PepSurf. MimoTree, EpiSearch, and PepSurf were run individually to predict the epitope for each input mimotope set and its corresponding antigen structure using default parameters as described above. The prediction from MimoTree and the highest-ranked prediction from EpiSearch and PepSurf, respectively, were concatenated into a list, which is sorted from highest to lowest by the number of occurrences of the different elements in it. The residues in the predictions were weighted by the number of times they appear in this list. Since three approaches are here being combined, the largest number of occurrences is three, while the lowest number of occurrences is one. According to Mayrose et al. (Mayrose et al., 2007) and Huang et al. (Huang et al., 2008b), 95% of all available epitopes in the PDB have a solvent-accessible surface area of no more than 2000 Å2, and an epitope typically contains no more than 40 amino acids. In line with these observations, we took the top 45 residues from the prediction list combining the three methods as the prediction for the “The Ensemble Method” with MimoTree incorporated, effectively utilizing a majority voting scheme. See Fig. 2 for an overview of the approach.

**Figure 2.**
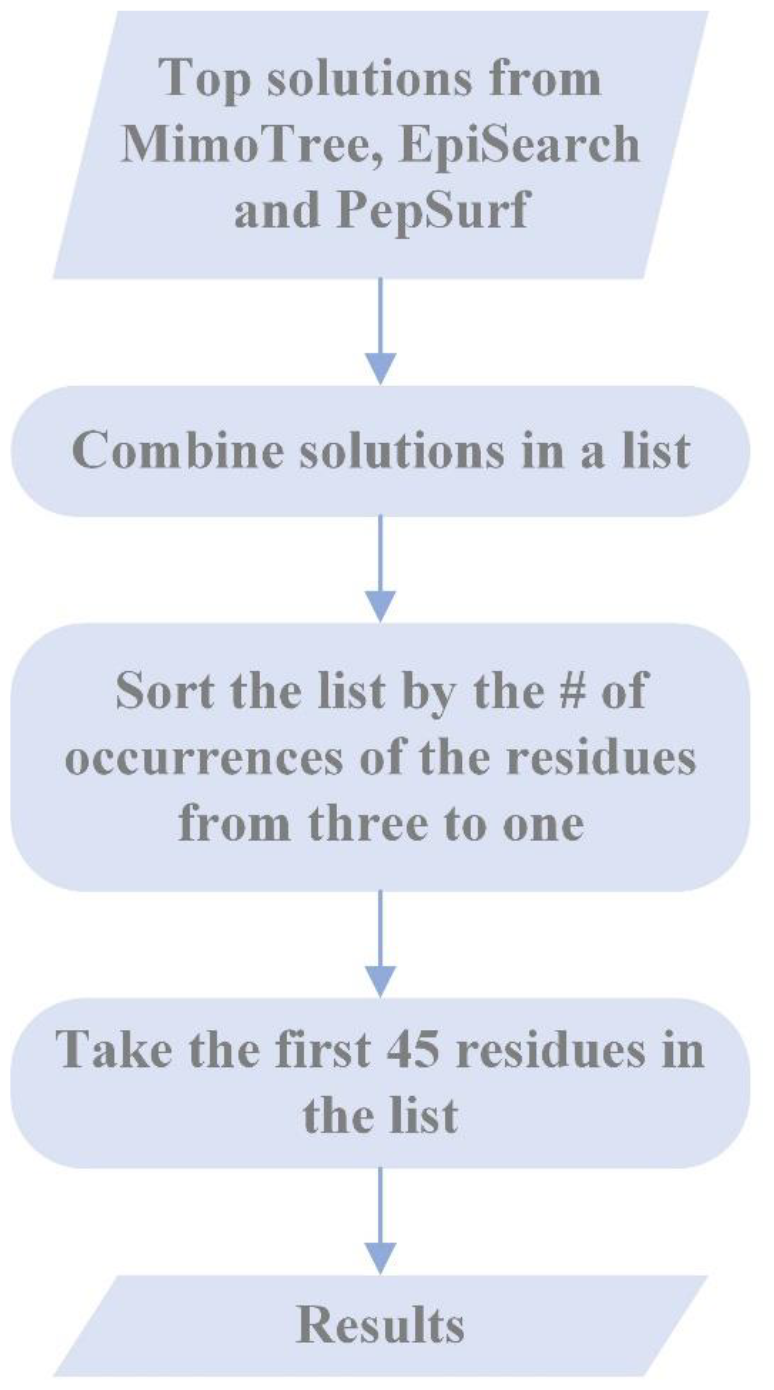
Flowchart of the incorporation of MimoTree into “The Ensemble Method”.

### 2.8 Statistical Analysis for Each Solution

To statistically evaluate the performance of the proposed methods, we use sensitivity, precision, Matthews Correlation Coefficient (MCC), and P-value on the predictions produced by the various methods on the test set.

#### 2.8.1 Sensitivity

Sensitivity is defined as the degree of coverage of the true epitope by the prediction. It is number residues correctly predicted divided by the number of residues in the true epitope or:

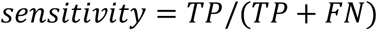

where TP is the number of true positives and FN is the number of false negatives in the prediction.

#### 2.8.2 Precision

Precision is the number of correctly predicted residues divided by the number of residues in the prediction. It is calculated as follows:

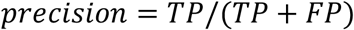

where FP is the number of residues in the prediction that are not in the true epitope.

#### 2.8.3 MCC

The Matthews Correlation Coefficient (MCC) is a coefficient of correlation between observed and predicted binary classifications. It is calculated as follows:

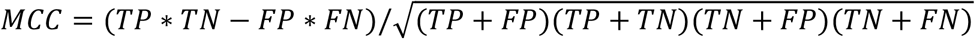

where TN is the number of residues in the antigen that are not part of the epitope.

#### 2.8.4 P-value

The P-value is defined as the probability that a random prediction for a given antigen can perform as well as or better than the prediction obtained by the various methods. The hypergeometric (HG) distribution describes the number of times that n objects of a specified type are successfully drawn (without replacement) from a finite number of N objects (containing M objects of the specified type). The probability of drawing k objects of the specified type is represented as follows:

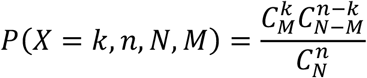

where k∈{1, 2, …, min (n, M)} and 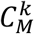 is determined as the number of combinations of M objects taken k at a time (Berkopec, 2007).

In evaluating the performance of the various methods across the test set, the P-value calculated based on the HG distribution represents the probability of randomly drawing n residues from an antigen containing N residues, of which M residues are in the prediction, and k or more residues are in the true epitope. A prediction with a P-value less than 0.05 is considered to be statistically significant. P-values are calculated as follows:

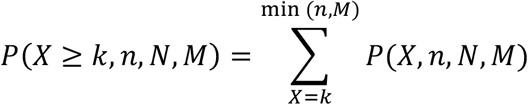

We also conducted tests of statistical significance to verify whether the performance differences observed between MimoTree and existing methods (EpiSearch and PepSurf) are statistically significant. First, the Shapiro-Wilk test was performed separately on the performance evaluations of MimoTree, EpiSearch, and PepSurf to confirm the normality of the data. Then, for the groups of data that are normally distributed, the 2-sided f-test was used to test whether the specific evaluations between MimoTree and PepSurf or EpiSearch, respectively, have equal variances. For the groups of data that have equal variances, the 2-sided t-test was performed to test the significance of the differences. For those with unequal variances, Welch’s t-test was performed to assess the significance.

## 3 Results

### 3.1 “The Ensemble Method”

First, we compared the performance of the “The Ensemble Method” to EpiSearch and PepSurf, respectively, across the test set containing ten antigen-antibody complex structures. The results are shown in Table 3. Overall, “The Ensemble Method” showed higher sensitivity, while largely maintaining the precision across the test set. The test case, 1YY9, is an exception in that no improvement was observed with “The Ensemble Method” since neither EpiSearch nor PepSurf were able to correctly predict any of the residues in the true epitope.

**Table 3.** Performance of “The Ensemble Method” Compared to EpiSearch and PepSurf.

| PDB ID | “The Ensemble Method” |  | EpiSearch |  | PepSurf |  |
| --- | --- | --- | --- | --- | --- | --- |
|  | Sensitivity | Precision | Sensitivity | Precision | Sensitivity | Precision |
| 1JRH | 1.00 | 0.38 | 0.21 | 0.13 | 0.79 | 0.44 |
| 1BJ1 | 0.31 | 0.39 | 0.17 | 0.33 | 0.25 | 0.45 |
| 1G9M | 0.38 | 0.14 | 0.00 | 0.00 | 0.38 | 0.17 |
| 1E6J | 0.93 | 0.33 | 0.93 | 0.62 | 0.00 | 0.00 |
| 1N8Z | 0.69 | 0.31 | 0.56 | 0.31 | 0.44 | 0.50 |
| 1IQD | 0.63 | 0.24 | 0.63 | 0.40 | 0.25 | 0.15 |
| 1YY9 | 0.00 | 0.00 | 0.00 | 0.00 | 0.00 | 0.00 |
| 2ADF | 0.56 | 0.36 | 0.13 | 0.13 | 0.56 | 0.75 |
| 1AVZ | 0.60 | 0.23 | 0.00 | 0.00 | 0.60 | 0.60 |
| 1HX1 | 0.65 | 0.31 | 0.55 | 0.44 | 0.10 | 0.10 |
| Average | 0.58 | 0.27 | 0.32 | 0.24 | 0.34 | 0.32 |

### 3.2 MimoTree

The performance of MimoTree was assessed on the same test set, and the results are given in Table 4.

**Table 4.**
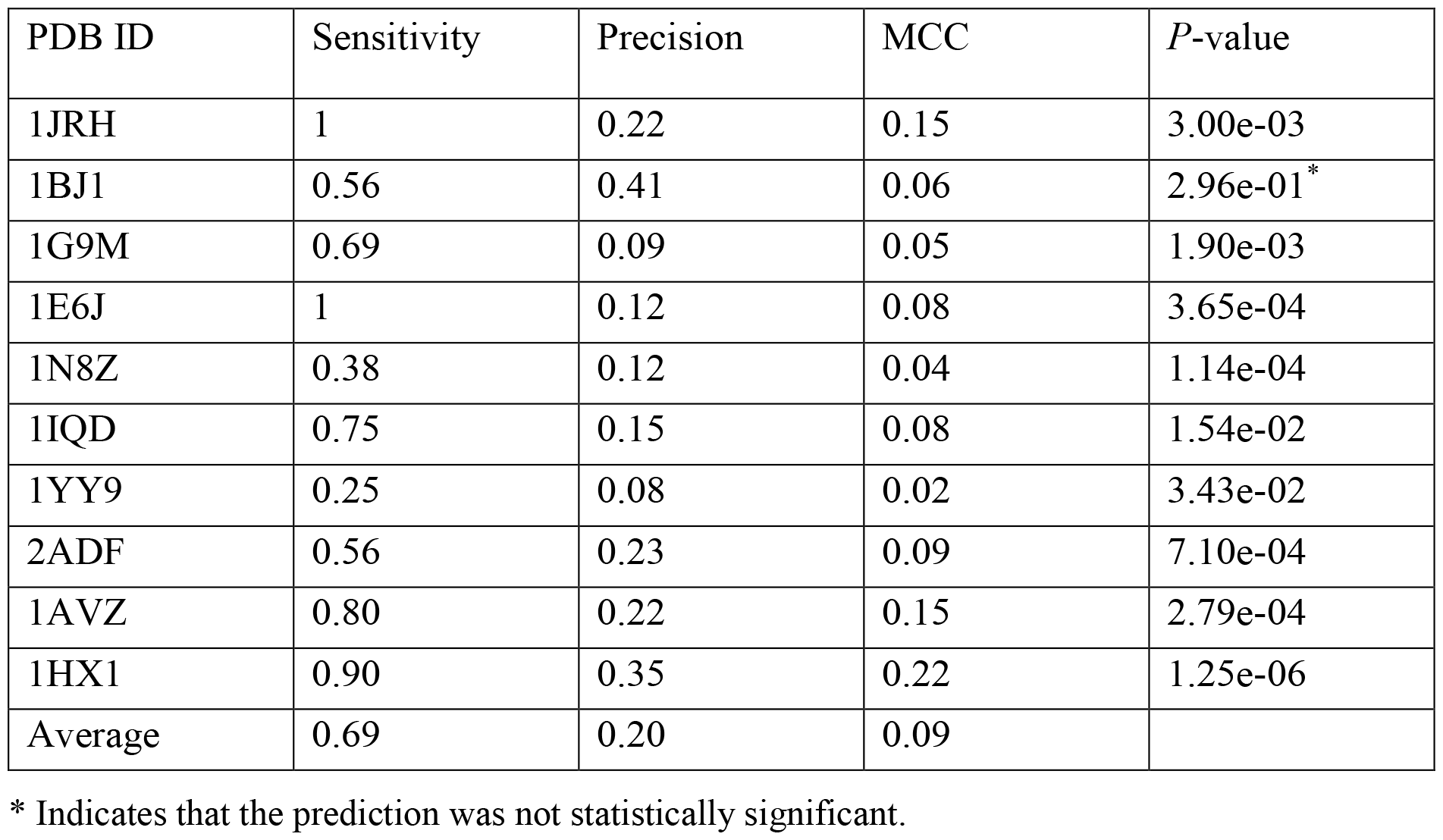
Performance of MimoTree Across the Test Set.

We further evaluated the performance of MimoTree using antigen-only structures as input. The results are given in Table 5. As expected, the sensitivity is somewhat diminished compared to when the antigen-antibody complex structures are used. None of the other published methods have been evaluated on antibody-only structures.

**Table 5.** Performance of MimoTree Using Unbound Antigen Structures as Input.

| PDB ID | Sensitivity | Precision | MCC | *P*-value |
| --- | --- | --- | --- | --- |
| 1VPF | 0.53 | 0.39 | 0.06 | 0.52e-01* |
| 3DNO | 0.38 | 0.16 | 0.05 | 0.83e-02 |
| 1A8O | 1.00 | 0.11 | 0.07 | 3.52e-04 |
| 1D7P | 0.50 | 0.11 | 0.06 | 1.36e-02 |
| 1NQL | 0.00 | 0.00 | 0.00 | 2.15e-01* |
| 1AO3 | 0.44 | 0.21 | 0.09 | 6.91e-03 |
| 1AVV | 0.59 | 0.25 | 0.15 | 2.87e-03 |
| 1I6Z | 0.50 | 0.15 | 0.12 | 0.89e-06 |
| Average | 0.49 | 0.17 | 0.08 |  |

\* Indicates that the prediction was not statistically significant

### 3.3 Incorporation of MimoTree into “The Ensemble Method”

The performance of the incorporation of MimoTree into “The Ensemble Method” was also evaluated across the test set. The results are shown in Table 6.

**Table 6.**
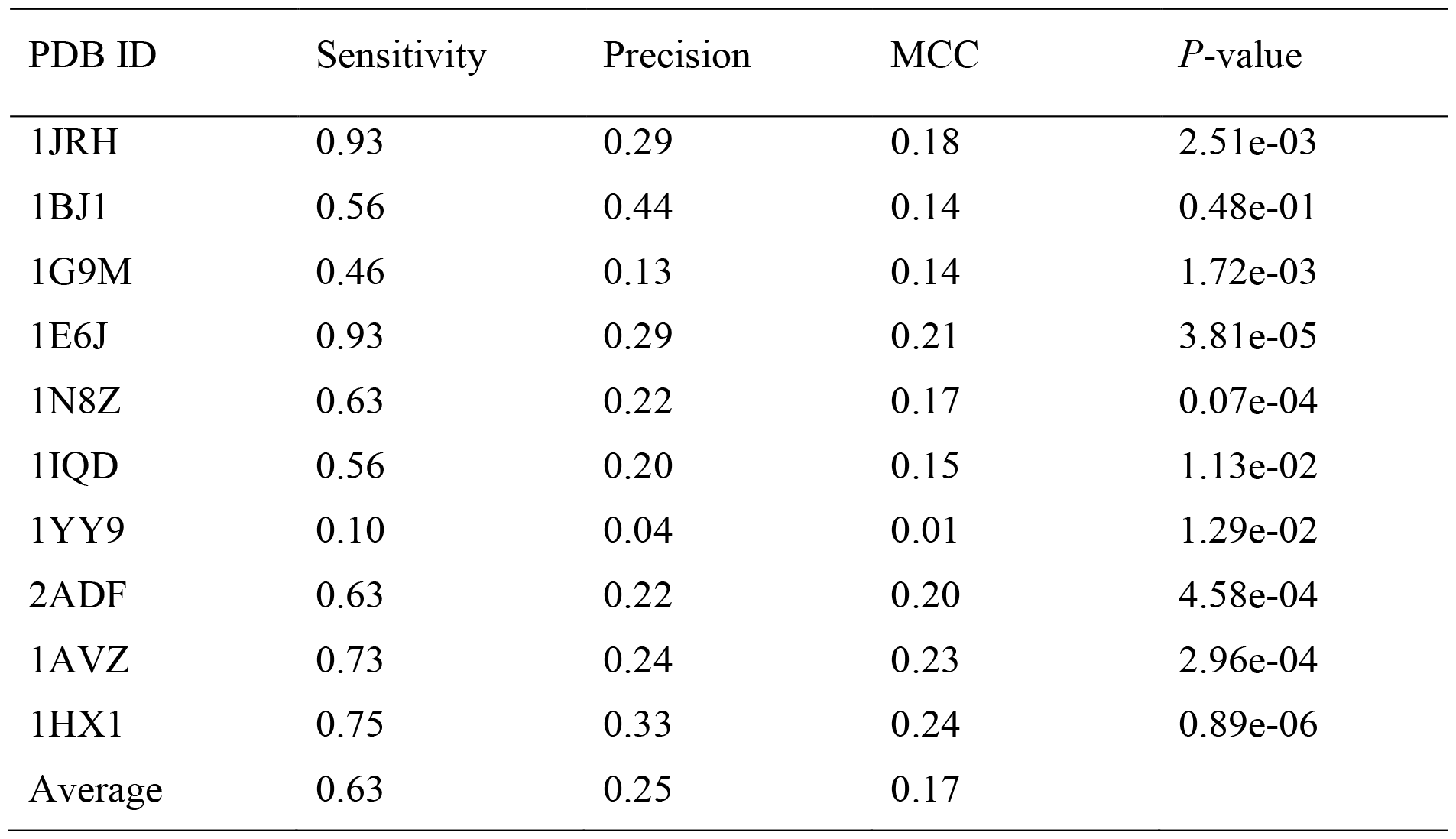
The Performance of “The Ensemble Method” Plus MimoTree Across the Test Set.

| PDB ID | Sensitivity | Precision | MCC | *P*-value |
| --- | --- | --- | --- | --- |
| 1JRH | 0.93 | 0.29 | 0.18 | 2.51e-03 |
| 1BJ1 | 0.56 | 0.44 | 0.14 | 0.48e-01 |
| 1G9M | 0.46 | 0.13 | 0.14 | 1.72e-03 |
| 1E6J | 0.93 | 0.29 | 0.21 | 3.81e-05 |
| 1N8Z | 0.63 | 0.22 | 0.17 | 0.07e-04 |
| 1IQD | 0.56 | 0.20 | 0.15 | 1.13e-02 |
| 1YY9 | 0.10 | 0.04 | 0.01 | 1.29e-02 |
| 2ADF | 0.63 | 0.22 | 0.20 | 4.58e-04 |
| 1AVZ | 0.73 | 0.24 | 0.23 | 2.96e-04 |
| 1HX1 | 0.75 | 0.33 | 0.24 | 0.89e-06 |
| Average | 0.63 | 0.25 | 0.17 |  |

### 3.4 Comparison with Existing Methods

A comparison of the performance of MimoTree, Ensemble + MimoTree, The Ensemble Method, EpiSearch, and PepSurf across the test set is shown in Table 7 and Fig. 3. Each group of sensitivity, precision, and MCC data for MimoTree, PepSurf, and EpiSearch, respectively, was found to be normally distributed through the Shapiro-Wilk test. The pairwise comparisons between the sensitivity, precision, and MCC for MimoTree and PepSurf or EpiSearch, respectively, were confirmed to have equal variances by a two-sided f-test, except for the precision of MimoTree vs. PepSurf. As such p-values for the comparisons were calculated by the two-sided t-test except for that one comparison which was by Welch’s t-test (see Table 8).

**Table 7.** Mean Sensitivity, Precision, and MCC by Approach.

|  | MimoTree | Ensemble +<br>MimoTree | Ensemble | EpiSearch | PepSurf |
| --- | --- | --- | --- | --- | --- |
| Mean sensitivity | 0.69 | 0.63 | 0.58 | 0.32 | 0.34 |
| Mean precision | 0.20 | 0.25 | 0.27 | 0.24 | 0.32 |
| Mean MCC | 0.09 | 0.17 | 0.10 | 0.08 | 0.05 |

**Table 8.** Statistical Significance Tests for Comparisons between MimoTree and Existing Methods.

|  | MimoTree vs. PepSurf | MimoTree vs. EpiSearch |
| --- | --- | --- |
| Sensitivity | 0.0044 | 0.0049 |
| Precision | 0.1153<sup>a*</sup> | 0.6173* |
| MCC | 0.9777* | 0.2110* |

<sup>a</sup> *P*-value calculated by Welch's *t*-test
\* Difference between the two groups of data is not statistically significant

**Figure 3.**
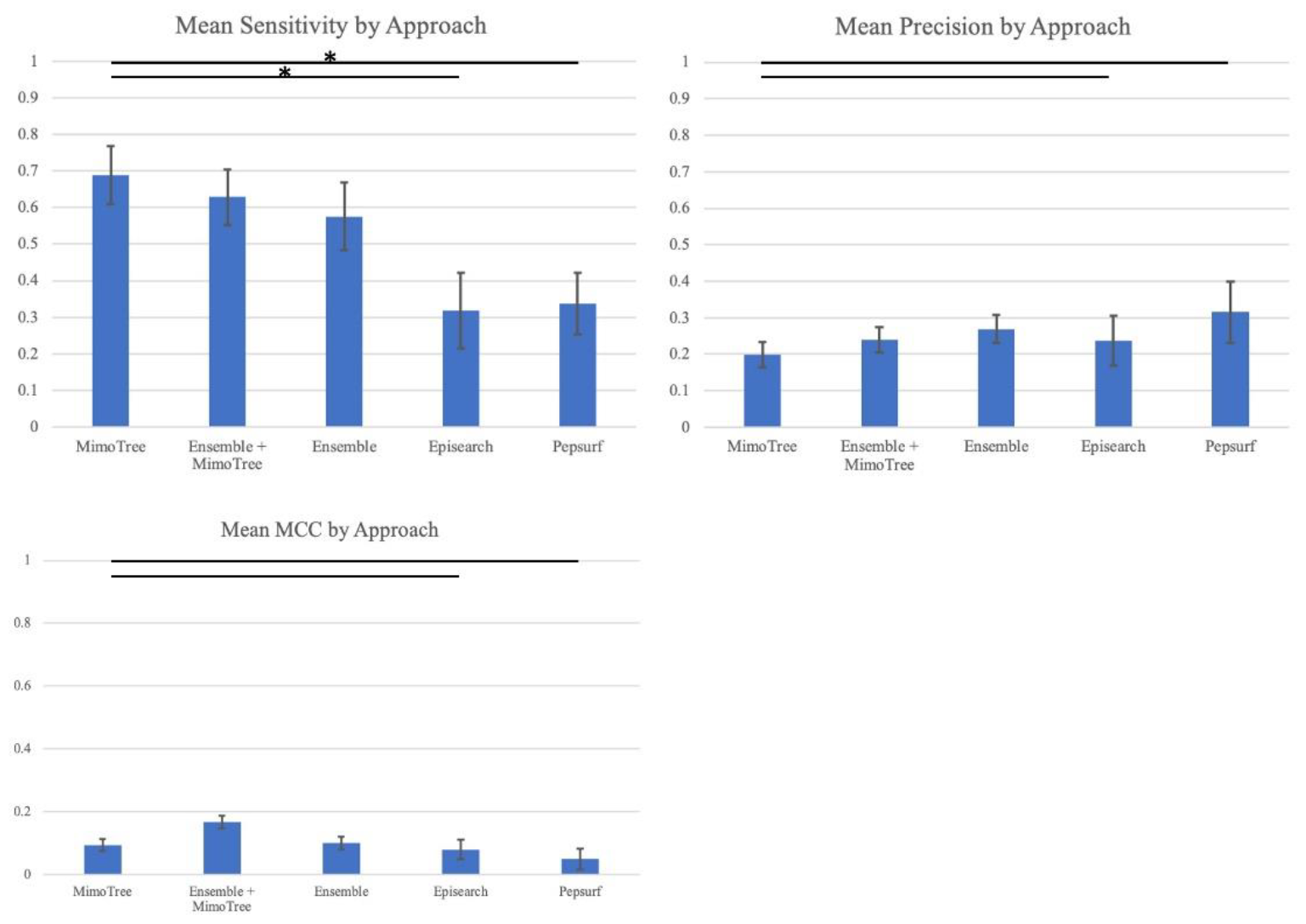
Bar plots of Mean Sensitivity, Precision, and MCC over ten case mimotope-to-epitope test set by approach with Standard Deviation indicated by the error bars. A * indicates difference is significant.

## 4 Discussion

In this study, we developed two novel methods for predicting the epitope on the antigen surface from mimotope data, “The Ensemble Method” and MimoTree. We also examined the effect of the incorporating MimoTree into the Ensemble approach. “The Ensemble Method” takes the union of the highest-ranked solutions obtained by EpiSearch and PepSurf. These two methods use different but complementary algorithms for the process of mapping mimotopes to an antigen structure, including differences in analyzing, scoring, and clustering the locations. EpiSearch first divides the surface of the antigen into overlapping surface patches. These patches are circular regions centered on each of the surface residues. EpiSearch then calculates the similarity between the surface residues contained in these circular patches and each input mimotope and obtains an initial epitope prediction. The manner by which EpiSearch divides the antigen surfaces tends to make this algorithm more suitable for predicting compact epitopes; if the true epitope is loosely distributed in multiple patches EpiSearch is less likely to predict it correctly as we see in the test set (e.g., for 1G9M, 1YY9, and 2ADF). Meanwhile, PepSurf uses color-coding techniques to find all possible linear paths on the antigen surface. These linear paths of different shapes may be able to traverse the entire antigen in one direction. Thus, these two methods are somewhat complementary in their surface searching approach. EpiSearch accurately locates specific residues in the epitope when it finds the right patches, but it also sometimes locates the wrong patches and therefore misses the correct epitope entirely. Conversely, PepSurf is better at finding the correct approximate epitope location but is less likely to find the specific residues within it. Our ensemble approach achieves a significantly higher sensitivity by taking the union of the highest-ranked solutions from both methods.

Our MimoTree algorithm is novel in that it addresses the possibility of gaps in mimotope-epitope mapping and it considers the 3D structure of the antigen in its final epitope connection step. If a mimotope cannot be continuously mapped on the antigen surface, several existing methods, including PepSurf, reflect this feature in the scoring of this path, i.e., a gap penalty with a negative value is added to the score. However, gaps in the mimotope sequenced matched to the epitope (as well as in the epitope itself if it is conformational) are often seen since mimotopes mimic the structure and sequence of epitopes, but are usually not the identical to the epitope. To solve this problem, MimoTree first unfolds the antigen surface into a surface map. For each input mimotope, MimoTree looks for each of its sequence fragments (seeds) on the antigen surface and tries to join these seeds in the sequence order of the original mimotope with a distance restraint based on the 3D structure of the antigen and the maximal length of the mimotope (if in an extended linear conformation). If the distance between the ends of the two seeds on the 3D antigen structure is less than or equal to the distance between corresponding residues in the mimotope (assuming a linear extended conformation), then the seeds are be connected. MimoTree predicts the mimotope to map to that position on the antigen surface. After obtaining the mappings on an antigen surface for the mimotope, MimoTree selects the top three seed connections in terms of overall length. The final output is the union of the three selected connections for each mimotope in the mimotope set. Because MimoTree uses a depth-first search method during the searching process, the longer the length of the seed connection (matching the mimotope to the antigen surface), the greater the degree of match overall.

While evaluating the performance of the existing methods, we found that EpiSearch and PepSurf were unable to accurately predict any of the residues in the epitope for the test case 1YY9_A. MimoTree, however, did find some matches between the mimotopes and the epitope for this test case. After carefully studying the antigen structure and the position of the epitope, we found that the residues in this epitope are loosely distributed and this antigen is a relatively large molecule (see Fig. 4), all of which greatly increases the difficulty of algorithmically predicting the location of the epitope. MimoTree has an advantage over the existing methods because it allows for some gaps between residues in the predicted epitope. Moreover, the other methods divide the antigen surface into several parts (e.g., patches with EpiSearch) and evaluate the similarity of these multiple antigen surface regions with the input mimotope sequence. Conversely, MimoTree directly uses mimotope sequence information to search the antigen surface for matching sequence fragments and attempts to concatenate these fragments. This more direct searching allows the algorithm to demonstrate better performance in predicting less-compact epitopes.

**Figure 4.**
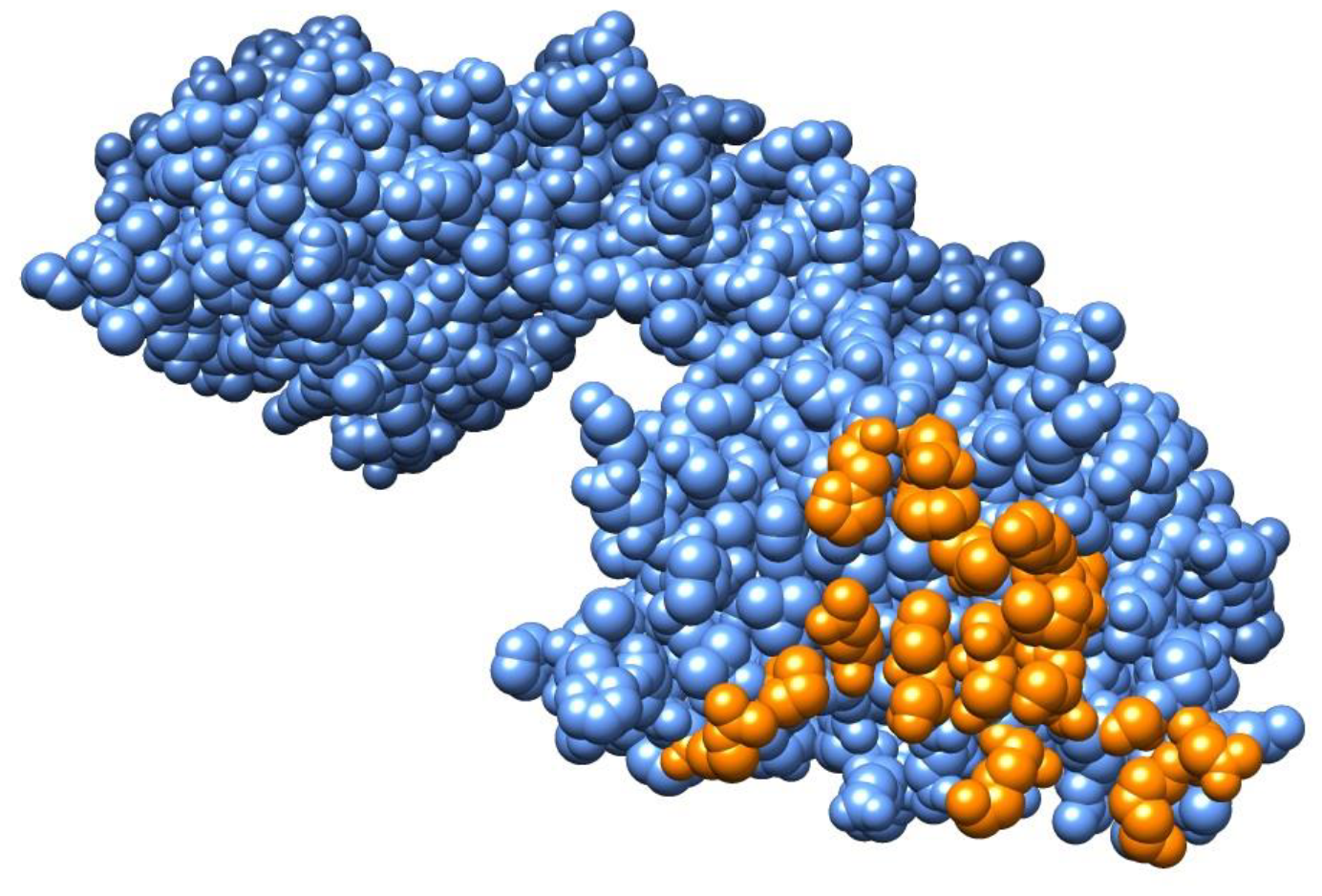
Structure of Epidermal Growth Factor Receptor (1YY9_A) with the true epitope shown in orange.

For 1HX1_B (bag chaperone regulator), the MimoTree prediction yielded a sensitivity of 0.90 and a precision of 0.35. Both sensitivity and precision are greatly improved compared to EpiSearch and PepSurf. After analysis, we found that the epitope of bag chaperone regulator is distributed widely across on the surface of the antigen (see Fig. 5). Since MimoTree analyzes the gaps formed in the alignments of mimotopes and the antigen, its search range is relatively wide. So, MimoTree performs better in this case where the epitope residues are widely distributed on the antigen surface.

**Figure 5.**
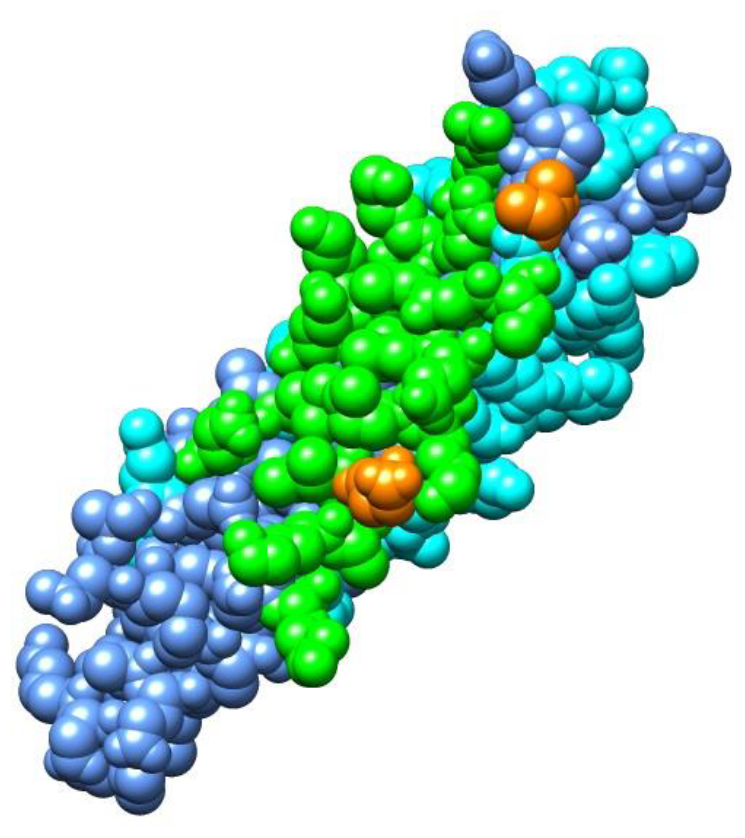
Structure of bag chaperone regulator (1HX1_B) showing the epitope predicted by MimoTree versus the true epitope. Residues that are in the true epitope and successfully predicted by MimoTree (true positives) are shown in green, those that are in the predicted epitope of MimoTree but not in the true epitope (false positives) are shown in cyan, and those that are in the true epitope but not predicted by MimoTree (false negatives) are shown in orange. The true epitope is in green and orange.

For 1BJ1_W, the only test case for which the prediction was not significant (having a p-value of 0.296, as given in Table 3), we found that the epitope consists of approximately 33% of the total number of residues in the antigen, which is a relatively large proportion. Therefore, it is more likely that randomly selected residues on the surface of this antigen will be part of the true epitope. Also, since MimoTree allows for the inclusion of gaps in mimotope-to-epitope mapping, the MimoTree predicted epitope may be more loosely distributed than the true epitope. While this often results in a prediction with higher sensitivity, it is also why the MimoTree prediction did not pass the hypothesis test in this case. It is an area for further optimization of the algorithm.

With respect to performance, there is usually a trade-off between sensitivity and precision. Because a larger number of residues are included in the prediction, the coverage of the epitope by the prediction is greater, thus increasing sensitivity. But the inclusion of more residues in the prediction may also mean that the proportion of correctly predicted residues decreases, resulting in decreased precision, and vice versa. This reasoning likely explains why incorporating MimoTree into the Ensemble approach did not improve the results.

In vaccine design, the use of inactivated virus tends to be avoided to limit side effects (Malonis et al., 2020). The use of mimotopes in vaccine design may be one way to minimize side effects. MimoTree maps sets of mimotopes to the antigen surface, which in turn enables the selection of mimotopes for vaccine development that correspond to epitopes in regions of the antigen structure with low frequencies of mutation to ensure evolutionary stability (Wang et al., 2021). If only the antigen structure is known (and not the antigen-antibody complex structure), selecting mimotopes in this manner is expected to limit the development of resistance to any resulting mimotope-based vaccine.

Since mimotopes are peptides with strong affinity to antibodies, when the antigen-antibody structure is known, MimoTree could still be used to enhance vaccine design because maximizing the degree of epitope coverage by the mimotopes may enhance the immunogenicity of the collection of mimotopes selected as raw material for vaccine production. This is likely a critical factor in vaccine design since the more complete the epitope coverage, the more effective the resulting mimotope-based vaccine may be in eliciting the desired immune response. Therefore, high sensitivity is expected to be more important for mimotope-to-epitope mapping for vaccine design.

The most promising raw materials for vaccines are those mimotopes that mimic the most antigenic regions (epitopes) and represent relatively evolutionarily stable epitope regions. Low precision may contaminate these evolutionary calculations through the inclusion of residues that are not part of the true epitope. In future iterations of MimoTree, it may be possible to enhance the precision by optimizing the combining of the predicted epitopes for each individual mimotope to yield the predicted epitope for the mimotope set by using some sort of voting rule instead of taking the union. In its current iteration, however, MimoTree outperforms the existing methods in terms of sensitivity without significantly compromising precision.

## 6 Author Contributions

R.L., M.S., A.C. and D.J-M. are responsible for the study conception and design. R.L. developed the algorithm and drafted the manuscript. S.W. contributed to data collection. S.W., M.S., D.V.E., A.C., and D.J.-M. contributed to the interpretation of the results and edited the manuscript. D.J.-M. oversaw the project. All authors contributed to the article and approved the submitted version.

